# The Polyadenosine RNA-Binding Protein ZC3H14 Localizes to Synapses and Regulates Synaptosomal CaMKIIα Levels

**DOI:** 10.1101/2020.10.08.331827

**Authors:** Stephanie K. Jones, Manushri Dalvi, Jennifer Rha, Sarah Kim, Kevin J. Morris, Omotola F. Omotade, Kenneth H. Moberg, Anita H. Corbett, Kenneth R. Myers

## Abstract

ZC3H14 (<u>Z</u>inc finger <u>C</u>ys<u>C</u>ys<u>C</u>ys<u>H</u>is domain-containing protein <u>14</u>), an evolutionarily conserved member of a class of tandem zinc finger (CCCH) polyadenosine (polyA) RNA binding proteins, is associated with a form of heritable, nonsyndromic autosomal recessive intellectual disability. Previous studies of a loss of function mouse model, *Zc3h14^Δex13/Δex13^*, provide evidence that ZC3H14 is essential for proper brain function, specifically for working memory. To expand on these findings, we analyzed the dendritic spines of hippocampal neurons from *Zc3h14^Δex13/Δex13^* mice. These studies reveal that loss of ZC3H14 does not affect dendritic spine density in either CA1 pyramidal neurons or dentate gyrus granule cells in the hippocampus. However, overexpression of ZC3H14 in cultured hippocampal neurons increases the overall density of spines. We next performed biochemical analyses of synaptosomes prepared from whole wild-type and *Zc3h14^Δex13/Δex13^* mouse brains to determine if there are changes in steady state levels of postsynaptic proteins upon the loss of ZC3H14. We found that ZC3H14 is present within synaptosomes and that a crucial postsynaptic protein, CaMKIIα, is significantly increased in these synaptosomal fractions upon loss of ZC3H14. Together, these results demonstrate that ZC3H14 localizes to synapses, that its levels influence dendritic spine morphology, and that its loss dysregulates synaptic CaMKIIα levels, suggesting a potential role for ZC3H14 in synaptic function and plasticity.

## Introduction

Brain development requires the formation of complex neural circuits where individual neurons communicate with one another at sites of connections termed synapses (Sheng & Hoogenraad, 2007; Yuste, 2011). The majority of excitatory synapses are positioned at dendritic spines, small protrusions found on the dendritic branches of neurons (Bourne & Harris, 2008; Segal, 2005). These spines function as discrete biochemical signaling centers and are one of the primary sites of information processing in the brain (Adrian et al., 2014; Chen & Sabatini, 2012; Svoboda et al., 1996). During development, spines are generally thought to transition from thin filopodia through a maturation process that involves interim spine types into mature mushroom-shaped spines (Berry & Nedivi, 2017). These spines remain dynamic even in mature neurons to support synaptic plasticity (De Roo et al., 2008). The size of the spine head positively correlates with the number of glutamate receptors at the postsynaptic surface (Borczyk et al., 2019; Noguchi et al., 2005; Nusser et al., 1998). Thus, the morphology of dendritic spines is tightly linked to synaptic strength. Consequently, defects in dendrite spine morphology are associated with numerous neurological disorders, including fragile X syndrome, autism, and epilepsy (Fiala et al., 2002; Forrest et al., 2018; Newey et al., 2005; Phillips & Pozzo-Miller, 2015).

Because dendrites and axons extend far away from the neuronal cell body, tight regulation of gene expression is essential for normal neuronal development and synaptic plasticity. Once exported from the nucleus, mRNAs are transported to local sites for translation, allowing new proteins to be rapidly synthesized at or near individual synapses in response to stimuli such as synaptic activity (Besse & Ephrussi, 2008; Holt & Schuman, 2013). This translation in dendrites provides a mechanism for maintaining and modifying the local proteome that is more rapid and efficient than synthesizing proteins in the cell body and transporting them to a specific site (Glock et al., 2017). Numerous studies have directly linked synaptic plasticity to local translation and have identified specific subsets of mRNAs that localize preferentially to dendrites and dendritic spines (Cajigas et al., 2012; Huber et al., 2000; Kang & Schuman, 1996; Poon et al., 2006). A large number of RNA binding proteins mediate the many events that comprise post-transcriptional gene regulation, including events that occur in the nucleus prior to export to the cytoplasm (Scherrer, 2018), such as splicing, editing, and polyadenylation and the cytoplasmic events that contribute to local translation (Glock et al., 2017). Numerous studies have linked RNA binding proteins to synaptic plasticity, learning and memory, and neurological disease (Cooper et al., 2009; Kapur & Ackerman, 2018; Liu-Yesucevitz et al., 2011; Scheper et al., 2007; Wang et al., 2016).

Many of the RNA binding proteins linked to neurological disease play multiple roles in post-transcriptional regulation of gene expression (Thelen & Kye, 2020). ZC3H14 (Zinc Finger CCCH-Type Containing 14) is an evolutionarily conserved, ubiquitously expressed polyadenosine RNA-binding protein (Leung et al., 2009). Mutations in the *ZC3H14* gene cause an autosomal-recessive, non-syndromic form of intellectual disability (Al-Nabhani et al., 2018; Kelly et al., 2012; Pak et al., 2011). Studies examining the essential budding yeast orthologue, Nab2, have identified roles in regulating poly(A) tail length, RNA splicing, and mRNA decay (Anderson et al., 1993; Hector et al., 2002; Marfatia et al., 2003; Schmid et al., 2015; Soucek et al., 2016). The *Drosophila* orthologue of ZC3H14, Nab2, also plays a role in poly(A) tail length control (Kelly et al., 2016; Kelly et al., 2014; Pak et al., 2011). Studies in mammalian cells show that regulation of poly(A) tail length is a conserved function of ZC3H14 (Kelly et al., 2014). Loss of Nab2 function in yeast or flies is lethal (Anderson et al., 1993; Pak et al., 2011) and mutant flies exhibit defects in locomotor behavior as well as abnormal brain morphology (Kelly et al., 2016; Pak et al., 2011). Work exploiting the *Drosophila* system showed that Nab2 function is essential in neurons as phenotypes observed in flies lacking Nab2 can be rescued by neuronal-specific expression of Nab2 (Pak et al., 2011). These studies provide insight into why mutations in the ubiquitously expressed *ZC3H14* gene cause neurological deficits.

Multiple isoforms of ZC3H14 are produced in mammals through alternative splicing (See Fig. 2C) (Leung et al., 2009). All four ZC3H14 isoforms include the essential zinc finger RNA-binding domain (Kelly et al., 2010); however, isoforms 1-3 contain an N-terminal Proline-Tryptophan-Isoleucine (PWI)-like domain as well as a predicted nuclear localization signal, while isoform 4 contains an alternative first exon. ZC3H14 isoforms 1-3 are ubiquitously expressed, whereas isoform 4 is expressed primarily in testis (Leung et al., 2009). While ZC3H14 isoforms 1-3 are primarily localized to nuclear speckles (Guthrie et al., 2011; Leung et al., 2009), analysis of cultured primary rat hippocampal neurons shows that ZC3H14 is found in both the nucleus and in neuronal processes (Bienkowski et al., 2017). Studies of cultured primary *Drosophila* neurons reveal that Nab2 is present within puncta in neurites, associated with both ribonucleoprotein complexes and polyribosomes (Bienkowski et al., 2017). In addition, cell fractionation assays from whole mouse brain reveal a cytoplasmic pool of ZC3H14, although the majority of the protein is found in the nucleus (Morris & Corbett, 2018). These observations regarding the localization of ZC3H14 are consistent with studies of the budding yeast Nab2 protein, which show that Nab2 is a shuttling RNA binding protein that can exit the nucleus in an poly(A) RNA-dependent manner (Green et al., 2002). This dynamic localization of Nab2/ZC3H14, means that ZC3H14 could regulate target RNAs in the nucleus and/or the cytoplasm.

To explore the function of ZC3H14 in mammals, a *Zc3h14* mutant mouse was generated (Rha et al., 2017). This mouse model removes exon 13 of *Zc3h14*, which is the first common exon present in all *Zc3h14* splice variants. This exon encodes part of the essential zinc finger RNA binding domain and thus no functional ZC3H14 protein is produced in homozygous *Zc3h14^Δex13/Δex13^* mice (Marfatia et al., 2003; Rha et al., 2017). These studies revealed that the ZC3H14 protein is not essential in mice; however *Zc3h14^Δex13/Δex13^* mice show defects in working memory, further supporting a role for ZC3H14 in normal brain function (Rha et al., 2017). Proteomic analysis comparing hippocampi from *Zc3h14^Δex13/Δex13^* mice to control *Zc3h14^+/+^* mice identified a number of proteomic changes that occur upon the loss of ZC3H14, including many changes in proteins with key synaptic functions. Mice lacking ZC3H14 show an increase in the steady-state levels of CaMKIIα, a protein kinase that plays a key role in learning and memory (Coultrap & Bayer, 2012; Coultrap et al., 2014; Herring & Nicoll, 2016; Kim & Hayashi, 2014; Rha et al., 2017). Furthermore, the ZC3H14 protein binds to *CaMKIIα* mRNA (Rha et al., 2017). Complementary work in *Drosophila* shows that Nab2 associates with *CaMKIIα* mRNA and represses a *CaMKIIα* translational reporter (Bienkowski et al., 2017). Taken together, these studies suggest a role for ZC3H14 in regulating expression of CaMKIIα. As CaMKIIα plays key roles in regulating dendritic spine morphology (Fukunaga et al., 2009), these results suggest that the loss of ZC3H14 could alter dendritic spine development; however, whether ZC3H14 is required for proper dendritic spine density or morphology has not yet been examined.

In this study, we analyze the *Zc3h14^Δex13/Δex13^* mouse model and find that loss of ZC3H14 does not affect dendritic spine density. However, overexpression of ZC3H14 increases the density of spines, in a ZC3H14 isoform-specific manner. Finally, ZC3H14 is present in synaptosomes and the loss of ZC3H14 leads to an increase in the steady-state level of CaMKIIα in synaptosomes. Our results demonstrate that ZC3H14 is found at synapses and that its overexpression affects dendritic spine density, suggesting a potential synaptic function.

## Material and Methods

### Mice

*Zc3h14^Δex13/Δex13^* mice were generated as previously described (Rha et al., 2017). To visualize dendritic spines in vivo, *Zc3h14^Δex13/+^* mice were crossed with the Thy1-GFP-M line reporter mice (The Jackson Laboratory, Stock #007788), which express enhanced GFP under the Thy1 promoter in a sparse subset of hippocampal neurons (Feng et al., 2000). Thy1-GFP-positive offspring heterozygous for the Zc3h14Δex13 allele were subsequently crossed with *Zc3h14^Δex13/+^* mice to generate *Zc3h14^+/+^* and *Zc3h14^Δex13/Δex13^* littermates carrying the Thy1-GFP transgene. Three-month-old *Zc3h14^+/+^*;Thy1-GFP and *Zc3h14^Δex13/Δex13^*;Thy1-GFP male littermates were used for *in situ* spine analyses. All procedures involving mice were performed in accordance with NIH guidelines for the use and care of live animals and were approved by the Emory University Institutional Animal Care and Use Committee.

### Primary hippocampal neuronal culture

Dissociated hippocampal neurons were cultured from embryonic day 17.5 (E17.5) mouse embryos of both sexes from a single litter. Briefly, brains were removed from embryos, and then hippocampi were dissected in ice cold Hank’s Balanced Salt Solution (HBSS). Hippocampi were pooled together and dissociated using trypsin for 12 minutes, then triturated and plated on 25 mm coverslips coated overnight with 100 μg/ml poly-L-lysine (Sigma). Neurons were plated at a density of approximately 350,000 cells per 35 mm dish. Cells were maintained in Neurobasal medium (ThermoFisher) supplemented with B-27 (ThermoFisher), penicillin/streptomycin, and GlutaMax (Invitrogen). Cells were fed once a week by replacing half of the medium with fresh growth medium.

### Transfection of hippocampal neurons

Hippocampal neurons were transfected at 11 days in vitro (DIV 11) using Calphos calcium phosphate transfection reagent (Takara) according to manufacturer’s protocol. The following DNA plasmids were used: GFP-ZC3H14-Isoform 1 and GFP-ZC3H14-Isoform 3 (Leung et al., 2009), LifeAct-mRuby (Riedl et al., 2008) (provided by Dr. Gary Bassell, Emory University), and eGFP-N1 (GFP). All DNA constructs were prepared using Endotoxin-free Maxi Prep kits (Qiagen).

### Immunofluorescence microscopy

Hippocampal neurons were fixed at either DIV 12 or DIV 19 using freshly prepared 4% paraformaldehyde (PFA) and 4% sucrose in phosphate-buffered saline (PBS) for 15 minutes at room temperature. Neurons were washed in PBS, then blocked and permeabilized in PBS supplemented with 5% normal goat serum (NGS), and 0.2% Triton X-100 for 1 hour at room temperature. Cells were incubated with primary antibodies (rabbit anti-RFP (600-401-279; Rockland; 1:1000) and mouse anti-GFP (A-11120; Invitrogen; 1,2000) for 1 hour, then with Alexa Fluor (Alexa-488 or Alexa-546; 1:750; ThermoFisher Scientific) secondary antibodies for 1 hour, and finally mounted with Fluoromount-G (Southern Biotech). Samples were blinded prior to image acquisition. Dendritic arbors were imaged using an epifluorescent Nikon Eclipse Ti inverted microscope with a 20x objective (Plan Fluor, 0.5 NA). Analysis, including tracing and Sholl analysis, was carried out using the ImageJ Simple Neurite Tracer plugin (Longair et al., 2011). Dendritic spines were imaged as Z-stacks comprised of 21 optical sections (0.2 µm step-size) using a Nikon C2 laser-scanning confocal system with an inverted Nikon Ti2 microscope (60x Plan Apo objective, 1.4 NA). For display purposes, 2D images were generated from maximum intensity projections of Z-stacks. For image analysis, dendrite and spine volumes were reconstructed from Z-stacks using the Filament Tracer module in Imaris (Bitplane). Spine heads were manually seeded prior to spine volume reconstruction. Density and classification of dendritic spines were determined using automated measurements produced from the Filament Tracer module. For each neuron imaged, spines were analyzed from four separate secondary or tertiary dendritic branches (a minimum of 25 μm long) at least 50 μm away from the soma. Spines were classified as either stubby, mushroom, or thin, following parameters previously described (Swanger et al., 2011). Briefly, stubby spines are defined as having a length <1 µm and a head/neck width ratio <1.5; thin spines are defined as having a length 1-5 µm and a head/neck width ratio <1.5; mushroom spines are defined as having a length <5 µm and a head/neck width ratio >1.5.

For in situ spine analysis, mice were transcardially-perfused with 4% PFA in PBS. Brains were then post-fixed in 4% PFA for 48 hours at 4°C, embedded in optimal cutting temperature (OCT) compound, and cryosectioned sagittally at 40 μm thickness. Free-floating sections were immunostained to enhance GFP signal. Sections were permeabilized in 0.5% Triton X-100 in PBS for 30 minutes at room temperature, then blocked overnight at 4°C in 10% NGS and 1% bovine serum albumin (BSA) in PBS. Sections were incubated with rabbit anti-GFP (Invitrogen, #A11122; 1:1000) in 0.1% Triton X-100 and 1% BSA in PBS overnight at 4°C, then incubated with Alexa Fluor-488 goat anti-rabbit (ThermoFisher, #A-11008; 1:500) overnight at 4°C. Sections were mounted using DAPI-containing Fluoromount-G mounting medium (Southern Biotech). Labeled sections were imaged on a Nikon C2 laser-scanning confocal microscope. Secondary apical dendrites of CA1 pyramidal neurons were imaged as Z-stacks of 20 μm depth with 0.2 μm Z-steps using a Nikon C2 laser-scanning confocal system with an inverted Nikon Ti2 microscope (60x Plan Apo objective, 1.4 NA) at 2.5x digital zoom. Secondary dendrites of dentate gyrus granule cells were imaged as Z-stacks of 8 μm depth with 0.1 μm Z-steps at 4x digital zoom. All images were blinded prior to analysis and spine density and morphology were analyzed using Imaris as described above.

### Synaptosomal Fractionation

Mice were sacrificed and whole brains isolated at postnatal day 0 (P0). Brains were immediately frozen on dry ice, and stored at −80°C. We employed validated PCR primers to confirm the genotype of these P0 brains (Rha et al., 2017). These whole brains were homogenized in Thermo Scientific Syn-PER Synaptic Protein Extraction Reagent with EDTA-free protease inhibitor, using a Dounce tissue grinder [three (3) *Zc3h14*^+/+^, three (3) *Zc3h14*^Δex13/Δex13^]. Fractionation to obtain the homogenate (Hom), cytosol (Cyt), and synaptosome (Syn) fractions was performed as described in the Thermo Scientific Syn-PER Synaptic Protein Extraction Reagent protocol.

### Immunoblotting

Samples from the synaptosomal fractionation were resolved by SDS-PAGE and transferred to a 0.2 μm nitrocellulose membrane (Bio-Rad Laboratories). After blocking non-specific binding with 5% nonfat milk in 1X TBST (Tris-Buffered Saline, 0.1% Tween 20) solution, the membranes were incubated with primary antibody [anti-ZC3H14 (Leung et al., 2009), CaMKIIα (Thermo Fisher 13-7300), PSD-95 (Sigma-Aldrich MAB1598), Synaptophysin (Sigma-Aldrich MAB5258)], followed by incubation with the appropriate horseradish peroxidase (HRP)-conjugated secondary IgG antibody (Jackson ImmunoResearch). Total protein was visualized by Ponceau staining. Quantification of chemiluminescence and Ponceau staining was carried out in ImageJ, with significance calculated with an unpaired *t* test.

## Results

### The loss of ZC3H14 does not alter spine density in the mouse hippocampus

To assess whether the loss of ZC3H14 alters dendritic spine density in the hippocampus, we used the Thy1-GFP-M reporter line (Feng et al., 2000). ZC3H14 is expressed in CA1 pyramidal excitatory neurons and dentate gyrus granule cells in the hippocampus (Fig. 1A). Thy1-GFP mice express GFP in a limited subset of neurons within these regions of the hippocampus, which allows for the tracing of individual dendritic spines (Fig. 1A). We first examined the spines of CA1 pyramidal neurons from 3-month-old male mice (Fig. 1B). Quantification of dendritic spine density (Fig. 1B) revealed no statistically significant difference in spine density upon the loss of ZC3H14 (*p*>0.05 by unpaired *t-*test; n=20 neurons, 4 mice). Similarly, we found no difference in spine density in granule cells (*p*>0.05 by unpaired *t-*test; n=32 neurons, 4 mice) between *Zc3h14^Δex13/Δex13^*and *Zc3h14^+/+^* mice (Fig. 1C). Together, this suggests that the loss of ZC3H14 in mice does not affect dendritic spine density in the hippocampus.

**FIGURE 1.**
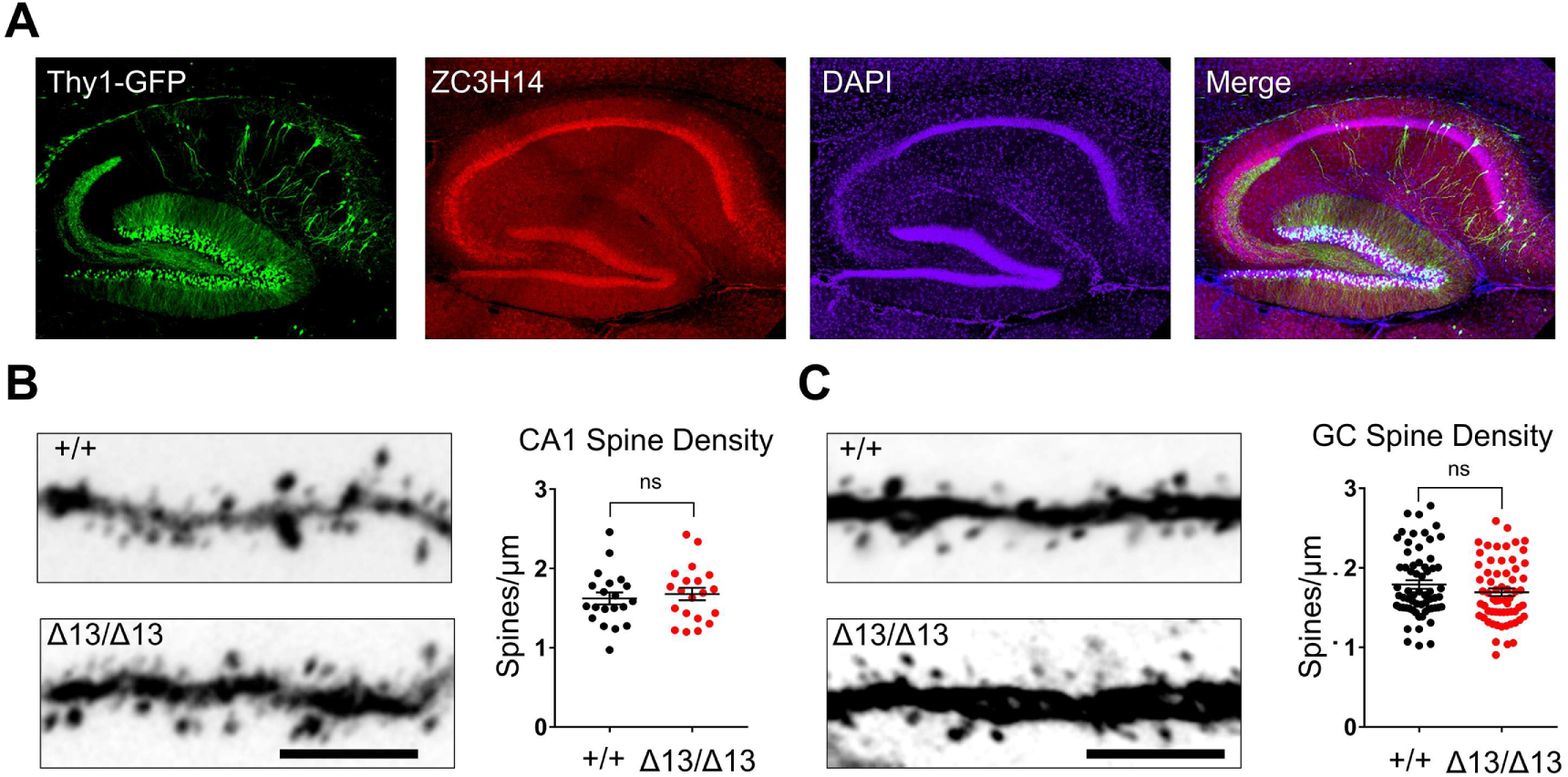
*Zc3h14^Δex13/Δex13^*mice display normal density of dendritic spines in the hippocampus relative to control mice. A) Representative images of hippocampal cryosections from Thy1-GFP (green) mice, stained for ZC3H14 (red) and DAPI (magenta). B) Left, representative images of spines from pyramidal CA1 neurons show no difference in spine density (n=20 neurons, 4 mice). Right, quantification of the number of dendritic spines per μm, comparing *+/+* and *Δ13/Δ13* mice with significance calculated by an unpaired *t* test (NS: *p*>0.05). C) Left, representative images of spines from dentate gyrus granule cells (n=32 neurons, 4 mice). Right, quantification of dendritic spines per μm, with significance calculated using an unpaired *t* test (NS: *p*>0.05).

### Overexpression of ZC3H14 increases dendritic spine density, specifically by increasing the number of thin spines

To examine whether ZC3H14 levels influence dendritic spine morphology, we examined whether overexpression of ZC3H14 impacts dendritic spine density in cultured hippocampal neurons. For this analysis, primary hippocampal neurons from control mice were co-transfected at DIV11 with Lifeact-mRuby, a small actin-binding peptide that labels dendritic spines (Riedl et al., 2008) to visualize dendritic spines, and either GFP (*+/+*), GFP-ZC3H14-Isoform1 (*+/+ (+Iso1)*), or GFP-ZC3H14-Isofom3 *(+/+ (+Iso3)*). We included two different isoforms of ZC3H14 (Fig. 2A,C) generated by alternative splicing (Leung et al., 2009). Both ZC3H14 Isoform 1 (Iso 1) and Isoform 3 (Iso 3) include the functionally important domains of ZC3H14, namely the zinc finger RNA binding domain (Fasken et al., 2019; Kelly et al., 2010) and the N-terminal PWI-like domain (Fasken et al., 2019; Grant et al., 2008), as well as a predicted nuclear localization signal (Fig. 2C). These ZC3H14 isoforms are primarily localized to the nucleus but can be detected in the cytoplasm of neurons (Bienkowski et al., 2017; Guthrie et al., 2011; Hector et al., 2002; Morris & Corbett, 2018).

**FIGURE 2.**
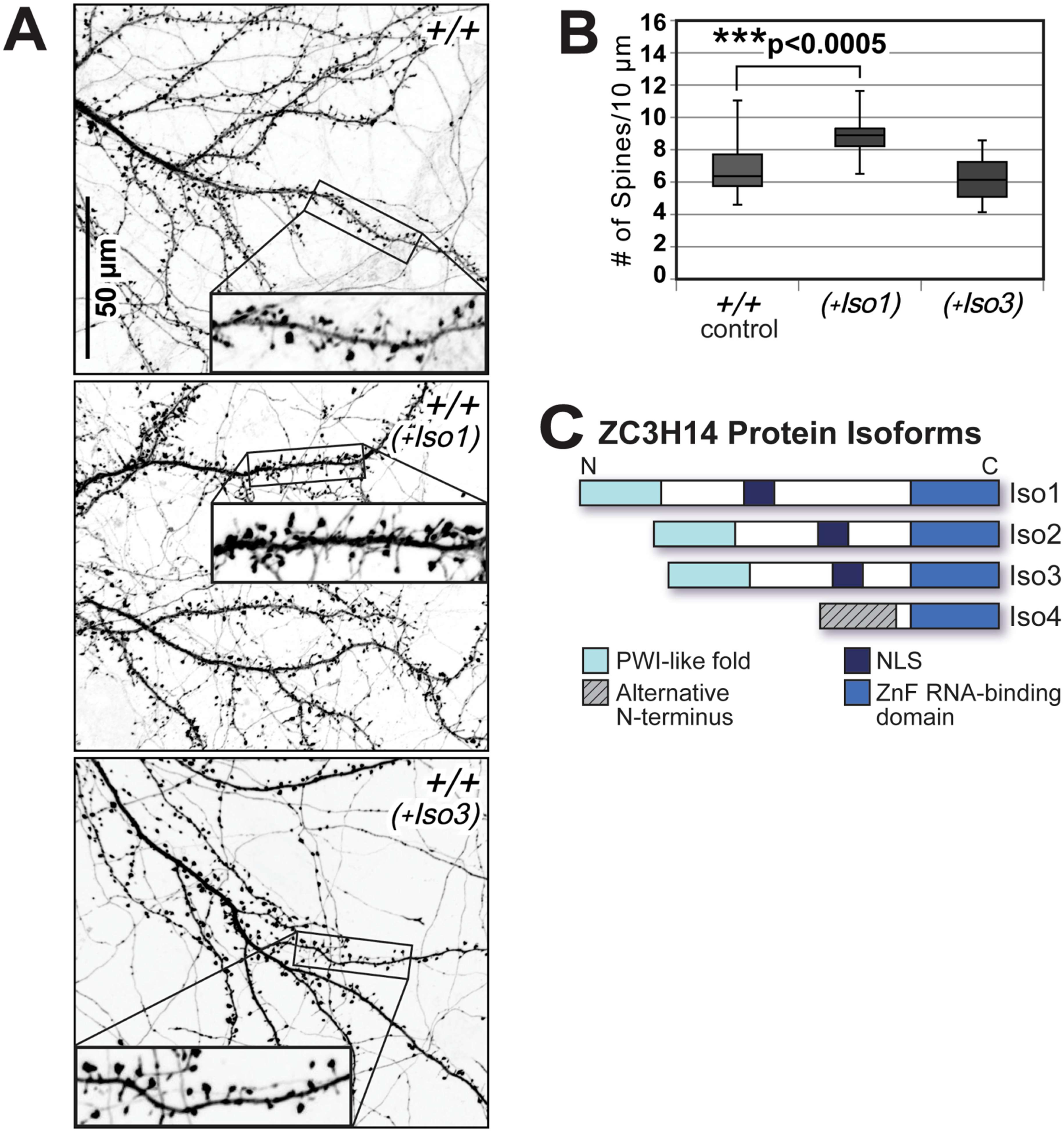
Overexpression of ZC3H14 Isoform 1 in cultured primary hippocampal neurons (DIV19) increases dendritic spine density. A) Representative inverted fluorescence images of control primary *Zc3h14^+/+^* (*+/+*) hippocampal neurons are shown together with *Zc3h14^+/+^* primary hippocampal neurons transfected with ZC3H14 Isoform 1 (*+/+ (+Iso1)*) or ZC3H14 Isoform 3 (*+/+ (+Iso3)*). The primary hippocampal neurons were cultured for 19 days *in vitro* (DIV19). Cultured neurons were fluorescently labeled by LifeAct-mRuby transfection. *Insets*, 2.4X magnification. B) Quantification of the number of dendritic spines per 10 μm, comparing *+/+* (n=21 neurons), *+/+ (+Iso1)* (n=14 neurons), and *+/+ (+Iso3)* (n=19 neurons) at DIV19 is shown. Statistical significance was calculated by an unpaired *t* test (NS>0.05; ***p<0.0005). C) A schematic of ZC3H14 protein isoforms 1-4 (Iso1-4) with labeled domains (Leung et al., 2009) is shown: Proline-Tryptophan-Isoleucine (PWI)-like fold domain, Alternative N-terminus, predicted Nuclear Localization Sequence (NLS), and CysCysCysHis (CCCH) zinc finger (ZnF) RNA-binding domain.

Neurons were then fixed and imaged at DIV19. Semi-automated quantification of spine density shows a statistically significant increase in spine density in *+/+ (+Iso1)* neurons as compared to control *+/+* [(*p*<0.0005; *+/+* n=21 neurons, *+/+ (+Iso1)* n=14 neurons)], but no significant difference in spine density in *+/+ (+Iso3)* neurons as compared to control *+/+* [(*p*>0.05; *+/+ (+Iso3)* n=19 neurons)] (Fig. 2B). This analysis reveals that overexpression of ZC3H14 Isoform 1, but not ZC3H14 isoform 3, causes a statistically significant increase in dendritic spine density *in vitro*.

The morphology of dendritic spines is heterogeneous, often representing differences in their stability and function (Bosch & Hayashi, 2012; Bourne & Harris, 2008). To determine if the overexpression of ZC3H14 affects spine morphology in DIV19 hippocampal neurons, we classified each spine as “Stubby”, “Thin”, or “Mushroom” (Arellano et al., 2007; Bourne & Harris, 2007; Peters & Kaiserman-Abramof, 1970; Swanger et al., 2011) according to both length and the head width to neck width ratio (Fig. 3A).

**FIGURE 3.**
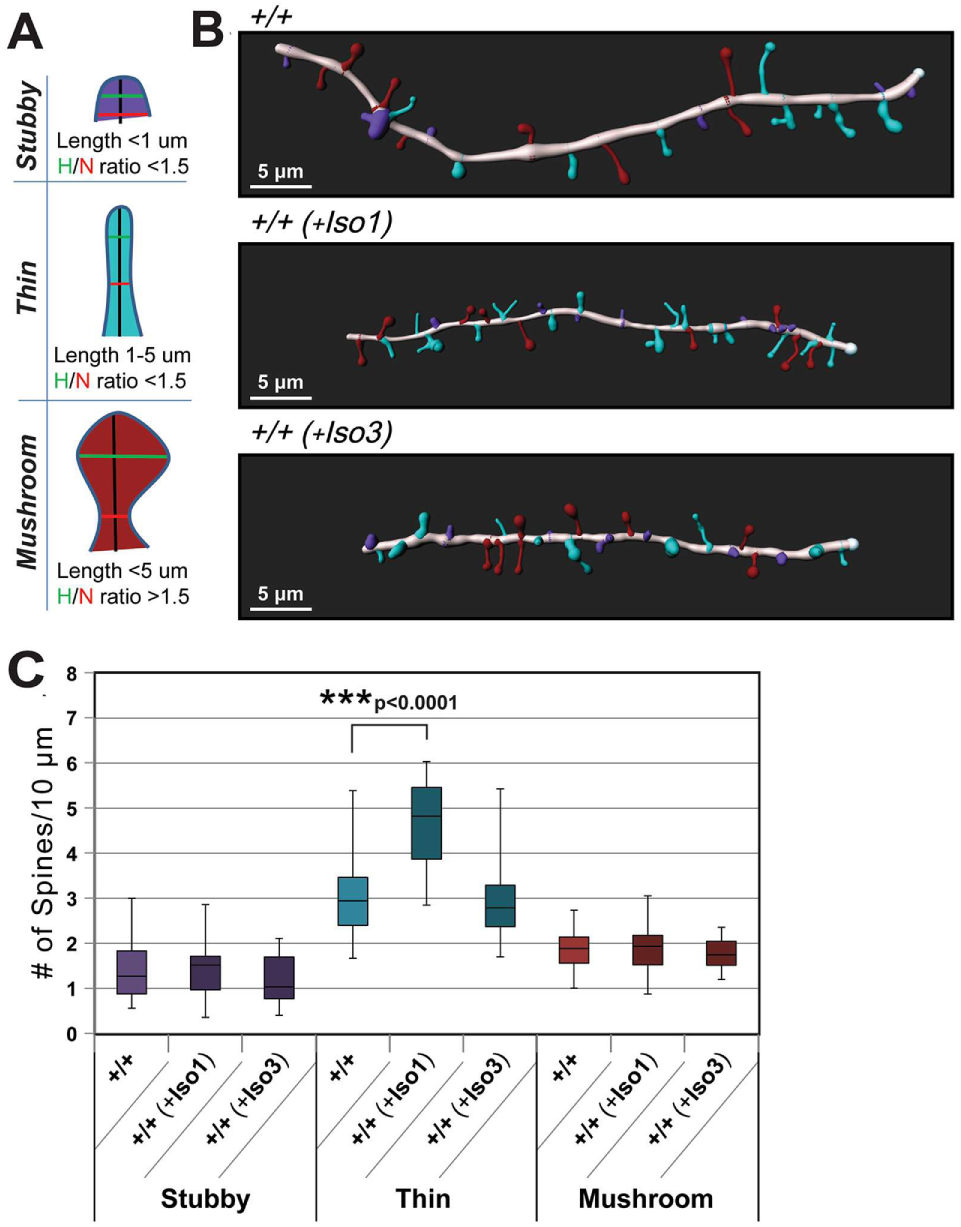
The increase in spine density detected in DIV19 primary hippocampal neurons that overexpress ZC3H14 Isoform 1 is due to an increase in the number of thin-type dendritic spines. A) A schematic illustrating the measurements used to classify “Stubby” (purple), “Thin” (light blue), and “Mushroom” (dark red) type dendritic spines, in terms of spine length, head width (H, green), and neck width (N, red). B) Representative Imaris software reconstructions of dendritic spines from *Zc3h14^+/+^*(*+/+*) primary hippocampal neurons (control), *Zc3h14^+/+^*neurons transfected with ZC3H14 Isoform 1 (*+/+ (+Iso1)*), and *Zc3h14^+/+^*neurons transfected with ZC3H14 Isoform 3 (*+/+ (+Iso3)*), were constructed from fluorescent images of 19 days *in vitro* (DIV19) cultured primary hippocampal neurons. C) Quantification of the number of each spine type per 10 μm, comparing *+/+* (n=21 neurons), *+/+ (+Iso1)* (n=14 neurons), and *+/+ (+Iso3)* (n=19 neurons) samples. Statistical significance was calculated by an unpaired *t* test (NS>0.05; ***p<0.001).

To assess whether overexpression of ZC3H14 alters the density of specific dendritic spine subtypes, 3D reconstructions of dendritic branches were generated to measure spine morphology and classify each spine as stubby- (purple), thin- (light blue), or mushroom-type (dark red) (Fig. 3B). Representative reconstructions for *+/+, +/+ (+Iso1)*, and *+/+ (+Iso3)* DIV19 primary hippocampal neurons are shown in Figure 3B. Semi-automated analysis revealed a significant increase in thin-type spines in neurons overexpressing ZC3H14 Isoform 1 [(compare *+/+* and *+/+ (+Iso1)* neurons (*p*<0.0001)] with no statistically significant differences in stubby- or mushroom-type (Fig 3C).There were no significant changes in any of the dendritic spine classifications in neurons overexpressing ZC3H14 Isoform 3 compared to *+/+* (Fig 3C). The increase specifically in thin-type spines, with no difference in stubby- or mushroom-type spines, in *+/+ (+Iso1)* DIV19 hippocampal neurons is consistent with a model where levels of ZC3H14 must be tightly regulated to ensure proper dendritic spine morphology.

### ZC3H14 is present in synaptosomes and loss of ZC3H14 results in increased steady state levels of CaMKIIα in synaptosomes

Previous work has demonstrated that there is a pool of ZC3H14 present in the cytoplasm of neurons (Bienkowski et al., 2017; Morris & Corbett, 2018). To determine whether ZC3H14 is present in synaptosomes, which contain postsynaptic dendritic spines and presynaptic terminals (Chicurel et al., 1993), fractionation was performed as illustrated in Figure 4A. We isolated three fractions from postnatal day 0 (P0) whole brains: the homogenate (Hom), cytosol (Cyt), and synaptosomes (Syn). Synaptophysin, a protein localized specifically to synaptic vesicle membranes (Wiedenmann & Franke, 1985), is enriched in the synaptosome (Syn) fraction from this preparation (Fig. 4B), providing evidence for successful enrichment of synaptosomal proteins. Immunoblotting performed on Hom, Cyt, and Syn fractions collected from *Zc3h14^+/+^* (*+/+*) and *Zc3h14^Δex13/Δex13^* (*Δ13/Δ13*) P0 whole brains show that ZC3H14 is detected in all three fractions, including synaptosomes (Fig. 4C). As a control, no full length ZC3H14 protein is detected in the samples from the *Δ13/Δ13* brains. This demonstrates that ZC3H14 is present in synaptosomes.

**FIGURE 4.**
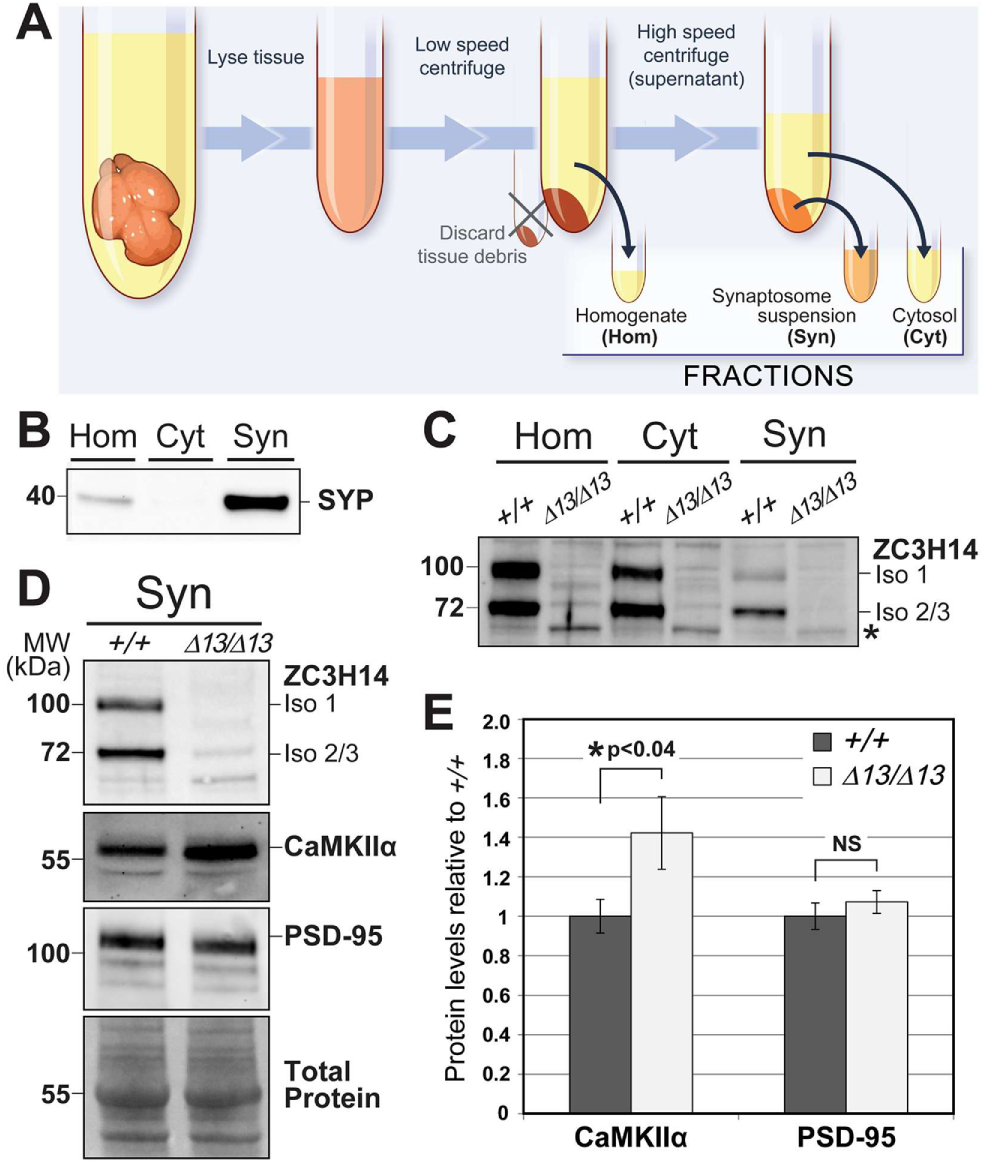
ZC3H14 is present in synaptosomes and CaMKIIα levels are increased in synaptosomal fractions from *Zc3h14^Δex13/Δex13^*mice compared to control. A) A schematic illustrates the synaptosomal fractionation procedure, which produces fractions corresponding to total homogenate (Hom), cytosol (Cyt) and synaptosome (Syn) derived from a single postnatal day 0 (P0) whole mouse brain. B) Immunoblotting for synaptophysin (SYP, ∼38 kDa), in the Hom, Cyt, and Syn fractions from a *Zc3h14^+/+^*(*+/+*) male whole mouse brain demonstrates the enrichment for this synaptic protein in the Syn fraction. C) An immunoblot analysis of Hom, Cyt, and Syn fractions comparing *+/+* and *Zc3h14^Δex13/Δex13^* (*Δ13/Δ13*) P0 whole mouse brain samples is shown. The top panel shows an immunoblot to detect ZC3H14, using an N-terminal antibody that detects ZC3H14 Isoforms 1 (∼100 kDa) and 2/3 (∼70 kDa) (Leung et al., 2009). The position of a truncated, nonfunctional ZC3H14 protein that is detected in the *Zc3h14^Δex13/Δex13^*mouse (Rha et al., 2017) is indicated by the *. D) Immunoblots shown compare the Syn fractions from *+/+* and *Δ13/Δ13* P0 whole mouse brain samples. Panels show immunoblots for ZC3H14, CaMKIIα (∼50kDa), and postsynaptic density protein 95 (PSD-95, ∼95 kDa). The bottom panel shows total protein detected by Ponceau, which serves as a loading control. E) Quantification of CaMKIIα and PSD-95 levels in the Syn fraction comparing *+/+* (n=3) and *Δ13/Δ13* (n=3) where n=independent male mice. For each protein, the level of protein detected was set to 1.0 for the *+/+* control and results are plotted as Relative to this control sample. Statistical significance was calculated by an unpaired *t* test (NS>0.05; \**p*=0.04).

As previous studies have implicated ZC3H14 in regulating the expression of CaMKIIα (Bienkowski et al., 2017; Rha et al., 2017), immunoblotting was performed to assess levels of CaMKIIα and PSD-95, a protein critical for maintaining the post synaptic density in dendritic spines (Hunt et al., 1996), specifically in synaptosomes (Fig. 4D). Results of this analysis show an increase in the level of CaMKIIα detected in the *Δ13/Δ13* synaptosomes compared to control, which is statistically significant (*p*<0.04; *+/+* n=3, *Δ13/Δ13* n=3) (Fig. 4E). In contrast, no statistically significant difference (*p*>0.05) was detected in the level of PSD-95 present in these synaptosomal fractions (Fig. 4D,E). Taken together, these data support a model where ZC3H14 is required for proper steady-state levels of CaMKIIα protein in synaptosomes.

## Discussion

This study employs a previously generated mouse model, *Zc3h14^Δex13/Δex13^* (Rha et al., 2017), to explore the requirement for ZC3H14 in dendritic spine density and morphology. Results of this analysis demonstrate that ZC3H14 is not required for proper dendritic spine density in the hippocampus. Studies in cultured primary hippocampal neurons reveal that overexpression of ZC3H14 increases overall dendritic spine density, primarily driven by an increase in the number of thin-type spines. As ZC3H14 is present in both the nucleus and synapses, these changes could result from altered post-transcriptional regulation of target transcripts. Finally, the loss of ZC3H14 causes an increase in the levels of CaMKIIα in synaptosomes, which could lead to defects in synaptic transmission or plasticity.

While the loss of ZC3H14 does not affect dendritic spine density, overexpression of ZC3H14 results in an increase in spine density, specifically the number of thin-type spines. Dendritic spines are highly heterogeneous in both function and morphology (Alvarez & Sabatini, 2007). Thin spines are fairly dynamic with long thin necks and small bulbous heads, which contain small excitatory synapses (Berry & Nedivi, 2017; Runge et al., 2020). Mushroom spines are more static with thin necks and broad heads that contain the largest excitatory synapses. Stubby spines are characterized by their lack of a definable head or neck, as well as their short squat appearance. While they contain large excitatory synapses, stubby spines are considered to be immature, and hence are not found as readily in adult brains (Harris et al., 1992). Previous studies have linked spine head volume with the size of the postsynaptic density (PSD), a dense and dynamic meshwork of proteins that mediate postsynaptic signaling (Arellano et al., 2007; Harris et al., 1992). Moreover, spine head volume correlates with AMPA receptor density and NMDA receptor-dependent calcium signaling, thus connecting spine morphology with synaptic strength (Majewska et al., 2000; Matsuzaki et al., 2001; Noguchi et al., 2005). The turnover of dendritic spines, as well as changes in their growth and retraction, is tied to alterations in brain circuitry that underlie learning and memory (Forrest et al., 2018; Yuste, 2011). For example, early longitudinal studies of the mouse barrel cortex demonstrated how spine and synapse formation/stabilization, as well as destabilization, occur in response to novel sensory experiences (Holtmaat et al., 2006; Zuo et al., 2005). Similarly, the size of individual spines and synapses from cultured neurons *in vitro* can be increased or decreased in response to specific patterns of activity (Bosch & Hayashi, 2012; Ho et al., 2011). For this reason, thin spines, which are more dynamic and structurally flexible, are thought to represent “learning spines” (Bourne & Harris, 2007), while mature mushroom-shaped spines likely represent “memory spines” (Borczyk et al., 2019; Bourne & Harris, 2007). Future experiments capable of monitoring spine dynamics would be required to determine whether overexpression of ZC3H14 affects spine motility or stability.

*Zc3h14^Δex13/Δex13^* mice display an approximately 40% increase in CaMKIIα levels in synaptosomes as compared to control *Zc3h14^+/+^* mice. CaMKIIα is a well-known regulator of synaptic plasticity and dendritic spine morphology (Coultrap & Bayer, 2012; Coultrap et al., 2014; Herring & Nicoll, 2016; Kim & Hayashi, 2014). We also identified ZC3H14 itself as a component of synaptosomes, raising the intriguing possibility that ZC3H14 could regulate the local translation of target transcripts, including *CaMKIIα* RNA. In a previous study characterizing the *Zc3h14^Δex13/Δex13^* mice, ZC3H14 was shown to bind the *CaMKIIα* transcript, and the loss of ZC3H14 was found to increase steady state levels of CaMKIIα in the brain (Rha et al., 2017). These findings are bolstered by studies in *Drosophila* in which the ZC3H14 orthologue Nab2 not only interacts with the *CaMKIIα* transcript but also represses a CaMKIIα translational reporter in neurons (Bienkowski et al., 2017). Previous studies have shown that *CaMKIIα* mRNA is localized to dendrites, and that CaMKIIα expression in dendrites is dynamically regulated by local translation (Burgin et al., 1990; Mayford et al., 1996; Paradies & Steward, 1997; Scheetz et al., 2000). Inhibiting CaMKIIα synthesis specifically in dendrites, but not in the soma, negatively affects synaptic plasticity and long-term memory but not learning (Aakalu et al., 2001; Ashraf et al., 2006; Miller et al., 2002; Neant-Fery et al., 2012). This is consistent with the working memory deficits observed in *Zc3h14^Δex13/Δex13^* mice (Rha et al., 2017) and with pan-neuronal depletion of Drosophila Nab2, which causes short-term memory deficits while leaving learning intact (Kelly et al., 2016).

The elevation of synaptosomal CaMKIIα in *Zc3h14^Δex13/Δex13^* mice raises the possibility that dysregulated CaMKIIα expression underlies defects in synaptic plasticity. The synaptic accumulation and activation/phosphorylation state of CaMKIIα are known to differentially affect the insertion or removal of AMPARs from the synaptic membrane. For example, during LTP the activation of CaMKIIα promotes the stabilization and retention of GluA1-containing AMPA receptors through phosphorylation of targets including GluA1 S831, while LTD promotes distinct differential CaMKIIα signaling states that favor AMPA receptor internalization and synaptic removal, including phosphorylation of GluA1 S567 (Coultrap et al., 2014). Previous studies have observed robust protein synthesis in hippocampal neurons following stimulation with the group I mGluR agonist DHPG, which induces a well-characterized form of protein synthesis-dependent long-term depression (mGluR-LTD) (Huber et al., 2000). Notably, tetanic stimulation of hippocampal slices induces an approximately 30% increase in dendritic CaMKIIα levels (Ouyang et al., 1999), a magnitude comparable to the elevation we observe at baseline in *Zc3h14^Δex13/Δex13^* synaptosomes. This raises the intriguing possibility that chronically elevated CaMKIIα in the absence of ZC3H14 disrupts the dynamic range of CaMKIIα-dependent signaling required for bidirectional synaptic plasticity. Future studies examining synaptic plasticity in *Zc3h14^Δex13/Δex13^* mice will be required to test this model.

Taken together, the results presented here suggest that ZC3H14 is not required for proper dendritic spine density in the hippocampus, but it is localized to synapses where it may regulate the levels of important synaptic proteins including CaMKIIα. Growing evidence places the RNA binding protein ZC3H14 in a group with other RNA binding proteins, including FMRP, that are implicated in regulating local translation at the synapse (Thelen & Kye, 2020). While further work is required to define the molecular mechanism by which ZC3H14 contributes to synaptic function, these findings suggest that dysregulated protein expression upon the loss of ZC3H14 may contribute to defects in synaptic plasticity and neuronal dysfunction.

## Data Availability Statement

The data that support the findings of this study are openly available for sharing. There are no data deposited into any repository.

## Acknowledgements

Financial support as follows - NIH F31 HD070735 (JR), F31 NS092437 (OFO), NIH MH107305, AG054206 and GM058728 (AHC and KHM), and a Whitehall Foundation research grant to KRM (2023-08-069). Research reported in this publication was supported in part by the Emory University Integrated Cellular Imaging Microscopy Core of the Emory Neuroscience NINDS Core Facilities grant, 5P30NS055077. The content is solely the responsibility of the authors and does not necessarily reflect the official views of the National Institute of Health. We thank Drs. James Zheng, Shannon Gourley, Jinnah Hyder, and Paul Garcia for helpful discussions and advice as well as members of the Moberg, Zheng, and Corbett laboratories for their support.

